# Efficient analysis of toxicity and mechanism of food contaminants using network toxicology and molecular docking strategy: A Case Study of Aflatoxin B1

**DOI:** 10.1101/2024.01.10.574998

**Authors:** Zi-Yong Chu, Xue-Jiao Zi

## Abstract

The present study aims to promote network toxicology and molecular docking strategies for the efficient evaluation of the toxicity of food contaminants. With the example of liver injury induced by the food contaminant Aflatoxin B1, this study effectively investigated the putative toxicity of food contaminants and the potentially molecular mechanisms. Initially, we obtained a preliminary overview of the toxicity of Aflatoxin B1 by using the ProTox-II and ADMETlab 2.0 databases. Subsequently, it was possible to identify 156 potential targets associated with Aflatoxin B1 and liver injury by using the ChEMBL, SwissTargetPrediction, GeneCards and DisGeNET databases. These were further refined by the STRING 5.0 database and Cytoscape 3.9.0 software for 23 core targets, including AKT1, SRC and EGFR. Then, GO and KEGG pathway analyses performed by Metascape database indicated that the core targets of Aflatoxin B1-induced hepatotoxicity were mainly enriched in cancer-related signalling pathways. Speedy molecular docking using Quick Vina confirmed the strong binding energy between Aflatoxin B1 and the core targets. In summary, Aflatoxin B1 may induce liver injury by regulating cell proliferation, cell survival, cell growth, cellular immune responses, and cellular signalling cascade responses in hepatocytes. We have provided a theoretical basis for understanding the molecular mechanism of Aflatoxin B1 hepatotoxicity and for the prevention and treatment of Aflatoxin B1-induced cancers in food contaminants. Furthermore, our network toxicology and molecular docking methods also provide an effective method for the rapid evaluation of the toxicity of food contaminants, which effectively solves the cost and ethical problems associated with the use of experimental animals.

**Graphical Abstract:** 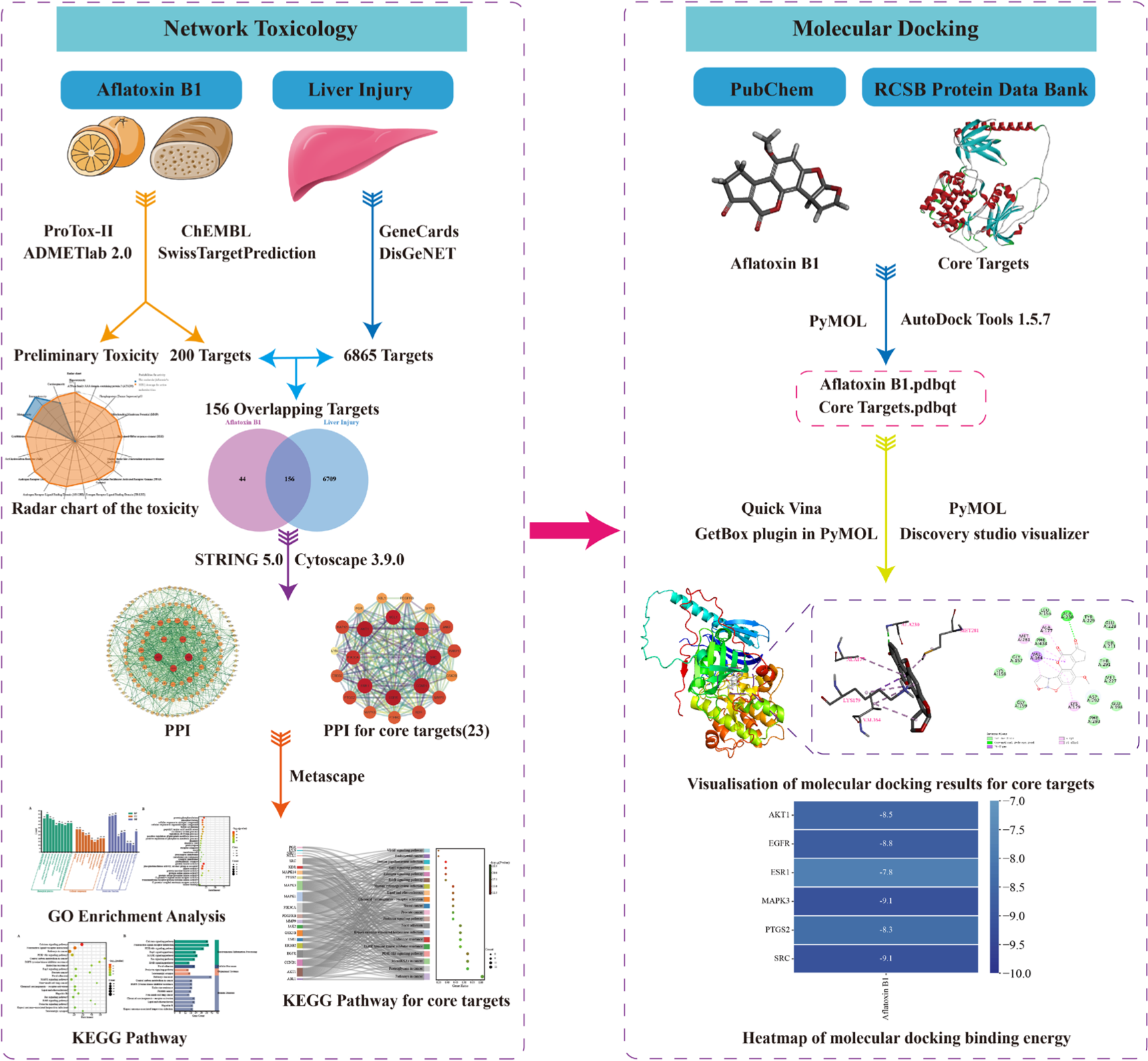

## 1. Introduction

Food safety is one of the most important issues of concern, as food quality and safety are vital to people’s health. Nevertheless, with increasing industrialisation and globalisation, the presence of food contaminants has become a focus of increasing concern[1]. The research on the toxicity of food contaminants is important for ensuring food safety, assessing potential risks in food and formulating appropriate food safety policies[2]. There are limitations in traditional toxicological assessment methods in the field of food safety in terms of time, cost, ethical issues of animal use and predictive power[3]. Therefore, there is a need for innovative approaches to comprehensively and effectively assess the potential health risks of an increasing number of uncharacterised toxic chemicals in the production, processing and preservation of food.

The combination of network toxicology and molecular docking for toxicity assessment of food contaminants is an efficient and accurate strategy. Network toxicology is a field of science that uses computer modelling and bioinformatics techniques to research the toxic effects of chemical substances on living organisms[4]. On the basis of constructing the interaction network between chemical substances and molecules in the organism, it predicts and evaluates the toxic effects of chemical substances on the human body, helping us to comprehend the mechanism of toxicity of chemical substances, screen for potentially toxic substances, and so on[5]. The molecular docking is a computational chemistry method used to predict the binding mode and affinity between small molecule compounds and target proteins[6]. In addition, the toxicity of a compound can also be determined based on the interaction and affinity between the compound and the target. The lower the ligand-receptor binding energy, the better the stability of the ligand-receptor binding conformation and the greater the potential for interaction[3]. On this occasion, we deployed network toxicology and molecular docking techniques to obtain insights into the potential toxicity and molecular mechanisms of Aflatoxin B1, a contaminant in food production, processing, and preservation.

Aflatoxin B1 is a natural toxin produced from *Aspergillus flavus* and *Aspergillus parasiticus*[7]. It is widely found in a variety of foods such as grains, nuts, legumes and spices[8]. Aflatoxin B1 has been recognised as a potent carcinogen and poses a potential hazard to human health. Prolonged exposure to Aflatoxin B1 may lead to a variety of health problems, including liver disease, liver cancer, immune system suppression and reproductive problems[9-11]. There are stringent regulatory standards and restrictions in place in many countries and regions to reduce the risk of Aflatoxin B1 to human health. The food safety agencies and agricultural departments usually monitor and control the level of Aflatoxin B1 in agricultural products to ensure food safety[12]. In summary, Aflatoxin B1 is a common naturally occurring toxin that poses a potential hazard to human health. Through monitoring, control and proper food handling practices, the risk of Aflatoxin B1 can be reduced.

Aflatoxin B1 has serious damaging effects on the liver[10]. Prolonged exposure to Aflatoxin B1 may lead to serious diseases such as hepatocellular damage, liver inflammation, liver fibrosis and liver cancer[10, 13]. Consequently, we implemented network toxicology and molecular docking strategies to rapidly investigate the toxic pathways involved in the hepatotoxicity of Aflatoxin B1. The purpose of our study was to gain insights into the toxic pathways involved in the hepatotoxicity of Aflatoxin B1, to clarify the toxicological profile of Aflatoxin B1, and to predicate its potential toxicity and molecular mechanisms by means of network toxicology and molecular docking techniques. Furthermore, this study may also provide insights into effective and rapid strategies for assessing the toxicity of food contaminants, as well as effectively addressing the limitations of traditional toxicological assessments in terms of time, cost, ethical issues of animal use, and predictive power, and establishing a basis for research to diagnose diseases associated with exposure to these toxic substances.

## 2. Methods

### 2.1. The preliminary network analysis of the toxicity of Aflatoxin B1

The structural information of Aflatoxin B1 was obtained from PubChem database, and then the structural information of Aflatoxin B1 was imported into ProTox-II and ADMETlab 2.0 to make a preliminary prediction of the toxicity of Aflatoxin B1 in accordance with the integration of the network search algorithm[14, 15].

### 2.2. Targets for collection of Aflatoxin B1

The structure of “Aflatoxin B1” was searched from PubChem database. The structure of Aflatoxin B1 was then entered into the ChEMBL database and SwissTargetPrediction database, and the species was restricted to “Homo sapiens”[16, 17]. Subsequently, potential targets of Aflatoxin B1 were retrieved from them. The results of our search were consolidated and de-duplicated, and the obtained target names were standardised using the Uniprot database[18]. These merged targets were then exploited to construct a library of Aflatoxin B1 targets.

### 2.3. Screening for hepatotoxicity-related targets

We searched the GeneCards and DisGeNET databases for relevant targets using the keywords “liver injury”, “hepatotoxicity” and “liver dysfunction”[19, 20]. In order to ensure that the obtained genes were highly correlated with hepatotoxicity and liver injury, the “score” threshold was set to the median, and the genes with “score” values greater than or equal to the median were selected and combined and de-weighted to establish a target gene library for hepatotoxicity[3]. In addition, we used the Venn diagram to screen for common potential targets between Aflatoxin B1 targets and liver injury sites, and regarded the intersection as a potential target of Aflatoxin B1 hepatotoxicity.

### 2.4. Protein interaction analysis and core target screening

The intersecting genes of potential targets of liver injury were entered into the STRING 5.0 database, restricting the species to Homo sapiens, and the interactions between the active target proteins corresponding to the target genes were analysed[21]. The results generated from the STRING 5.0 database were then imported into Cytoscape 3.9.0 to analyse and calculate the parameters of each node in the network graph, thus generating a protein-protein interaction (PPI) network graph[22]. Then, we used the following criteria to screen the core targets: the nodes corresponding to the targets satisfying the following conditions were selected as the core targets for Aflatoxin B1 hepatotoxicity: ① Closeness centrality > median, ② Radiality > median, and ③ Degree value > twice median. Finally, we used the MCODE plugin to validate the PPI network of these centrality genes[3].

### 2.5. Enrichment analysis of gene functions and pathways of target proteins

It was used to investigate the biological functions of potential targets in Aflatoxin B1-induced hepatotoxicity by collecting data for Gene Ontology (GO) analysis and Kyoto Encyclopedia of Genes and Genomes (KEGG) pathway enrichment analysis using the Metascape database[23]. First, we conducted GO analyses including biological process (BP), cellular component (CC), and molecular function (MF) assessments to elucidate their major biological functions[24]. Then, KEGG enrichment analysis was conducted to identify important pathways associated with potential targets of Aflatoxin B1-induced liver injury, and an FDR threshold of <0.05 was set to identify the major toxicity pathways of the obtained targets[25]. In addition, we also performed KEGG enrichment analysis of the core targets of Aflatoxin B1-induced hepatotoxicity using the Metascape database[23]. This analysis aimed to further investigate the significant signalling pathways involved in the core targets of liver injury. Finally, we performed visualisation analysis to effectively interpret and present the results of GO and KEGG analysis[26].

### 2.6. The rapid molecular docking of Aflatoxin B1 and core targets

To improve the efficiency and accuracy of molecular docking, we used Quick Vina for molecular docking to further analyse the intermolecular interactions between Aflatoxin B1 and the core target proteins of hepatotoxicity by predicting the binding mode and affinity[27]. First, We downloaded the 3D structure of Aflatoxin B1 from the PubChem compound database and saved it in sdf format. Next, We used ChemBioDraw 3D software to convert the sdf format to pdb format[28]. Then, we used AutoDock Tools 1.5.7 to pre-process the core target proteins downloaded from the RCSB Protein Data Bank (PDB) database and Aflatoxin B1 for molecular docking. Next, we used the GetBox plugin in PyMOL to determine the size of the molecular docking grid box. Then, we ran Quick Vina for molecular docking using the Anaconda Powershell Prompt command character. Finally, the results were visualised and analysed using Discovery studio visualizer and PyMOL and heat maps of molecular docking binding energies were drawn.

### 2.7. Statistical analysis

The data for the comprehensive study were obtained from an online database platform and were visualised and analysed using the SRplot online mapping platform[26].

## 3. Results

### 3.1. Toxicity assessment of Aflatoxin B1 in an initial network

Through integrating the results of toxicity analysis of Aflatoxin B1 by ProTox-II and ADMETlab 2.0, we obtained a basic summary of Aflatoxin B1 toxicity (**Fig. 1**). Toxicity modelling showed that the active toxicity endpoints were strongly associated with hepatotoxicity and liver injury. These discoveries are concordant with previous literary reports of Aflatoxin B1-mediated toxicity in humans, and provide a basis for further systematic and in-depth research on the toxic actions of Aflatoxin B1 in human beings[29].

**Fig. 1.**
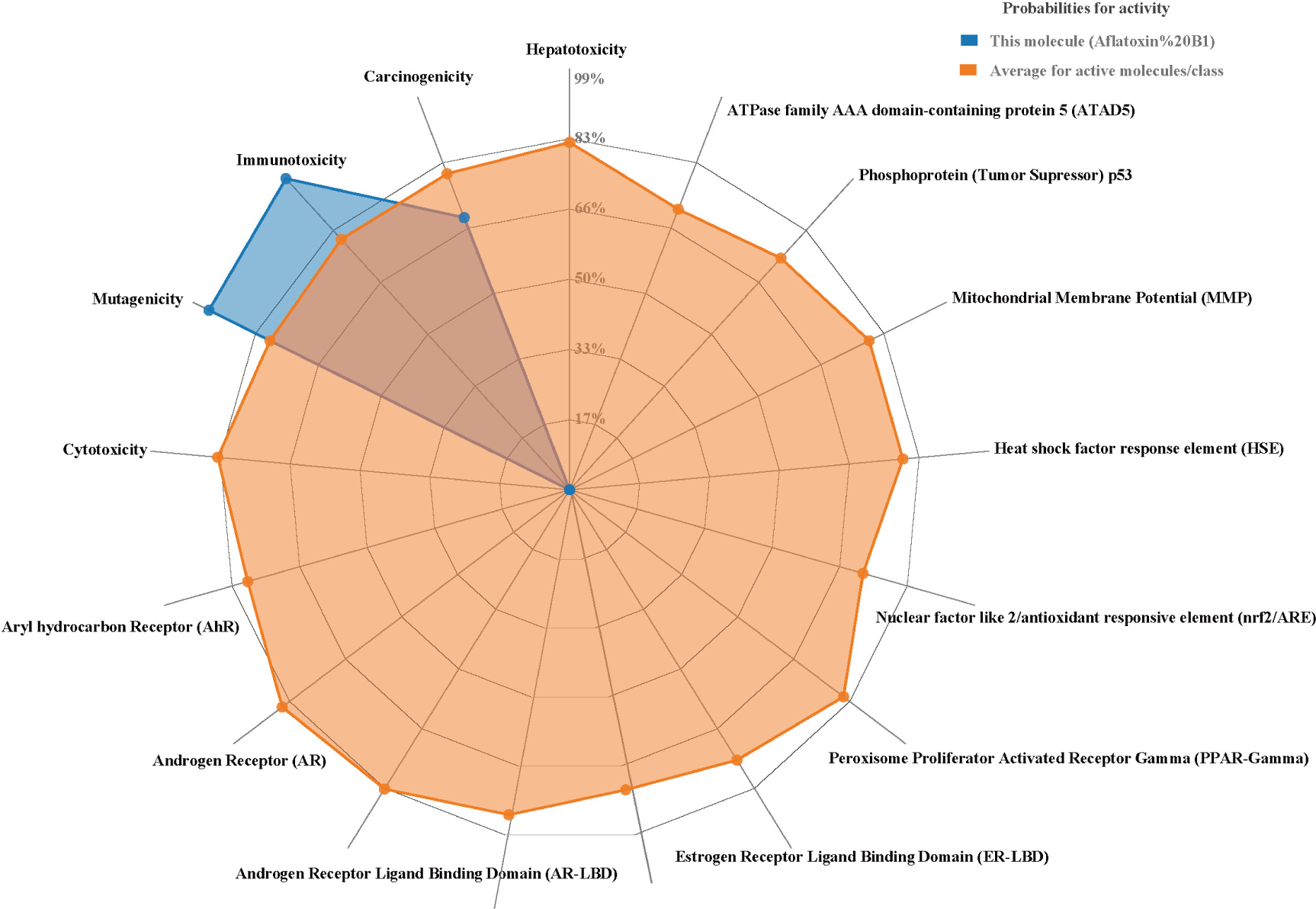
Radar chart of the toxicity of Aflatoxin B1.

### 3.2. Target of Aflatoxin B1-induced hepatotoxicity identified

At the present study, we initially screened 200 Aflatoxin B1 targets from the ChEMBL and SwissTargetPrediction databases. Then, we identified 6865 targets highly associated with hepatotoxicity through GeneCards and DisGeNET databases. The integration and deduplication of these target sets resulted in a total of 156 intersecting targets, which are potential targets for Aflatoxin B1-induced liver injury (**Fig. 2**).

**Fig. 2.**
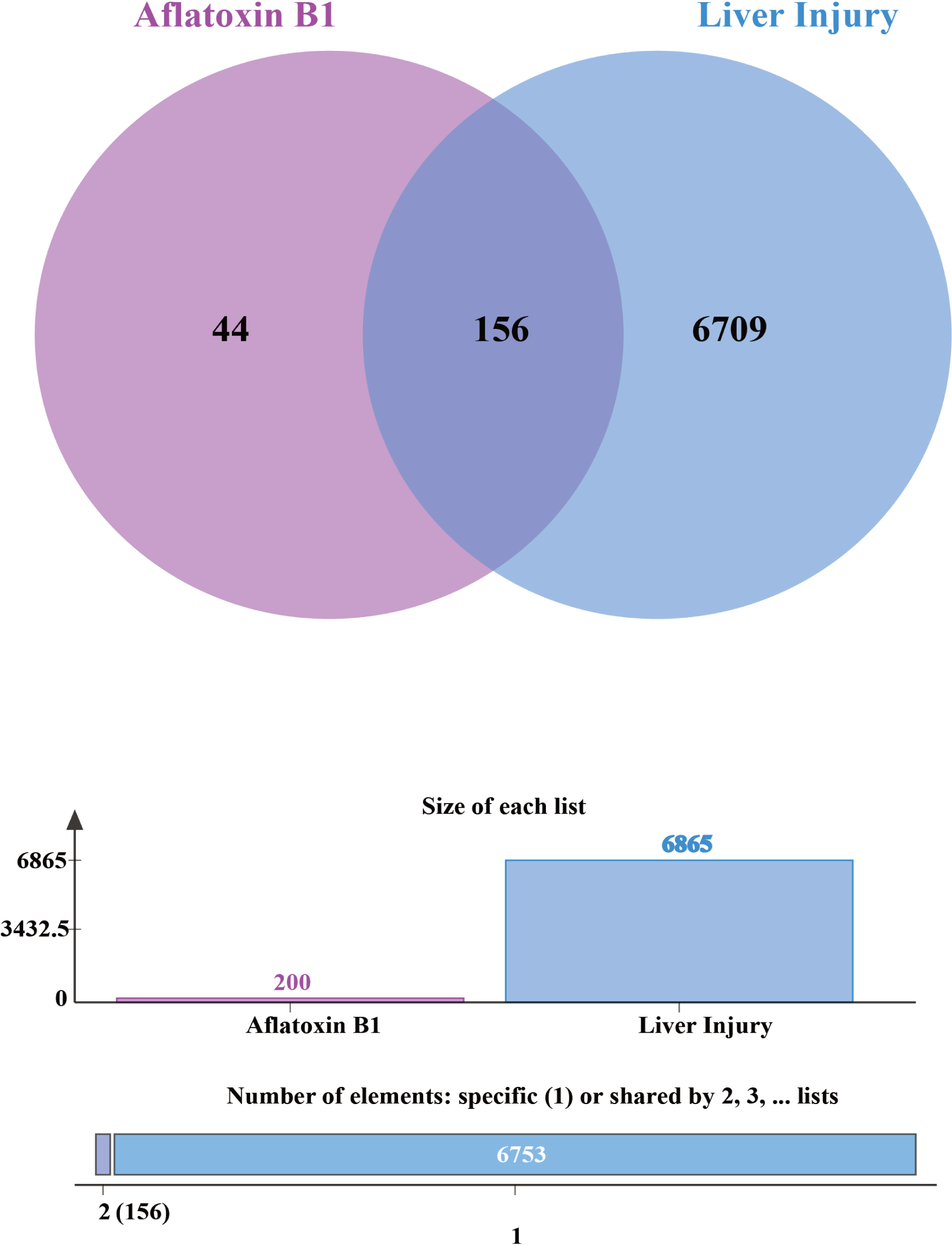
Venn diagram of Aflatoxin B1 and liver injury targets.

### 3.3. The potential target interaction network and the acquisition of core genes

We first constructed a protein-protein interaction network (PPI) with 154 nodes and 1705 edges with an average node degree value of 22.143 using the STRING 5.0 database. The topological properties of the network nodes, including Closeness Centrality, Degree, and Radiality, were analysed using Cytoscape 3.9.0 software. Meanwhile, we generated a visualised protein-protein interaction network diagram (**Fig. 3**).

**Fig. 3.**
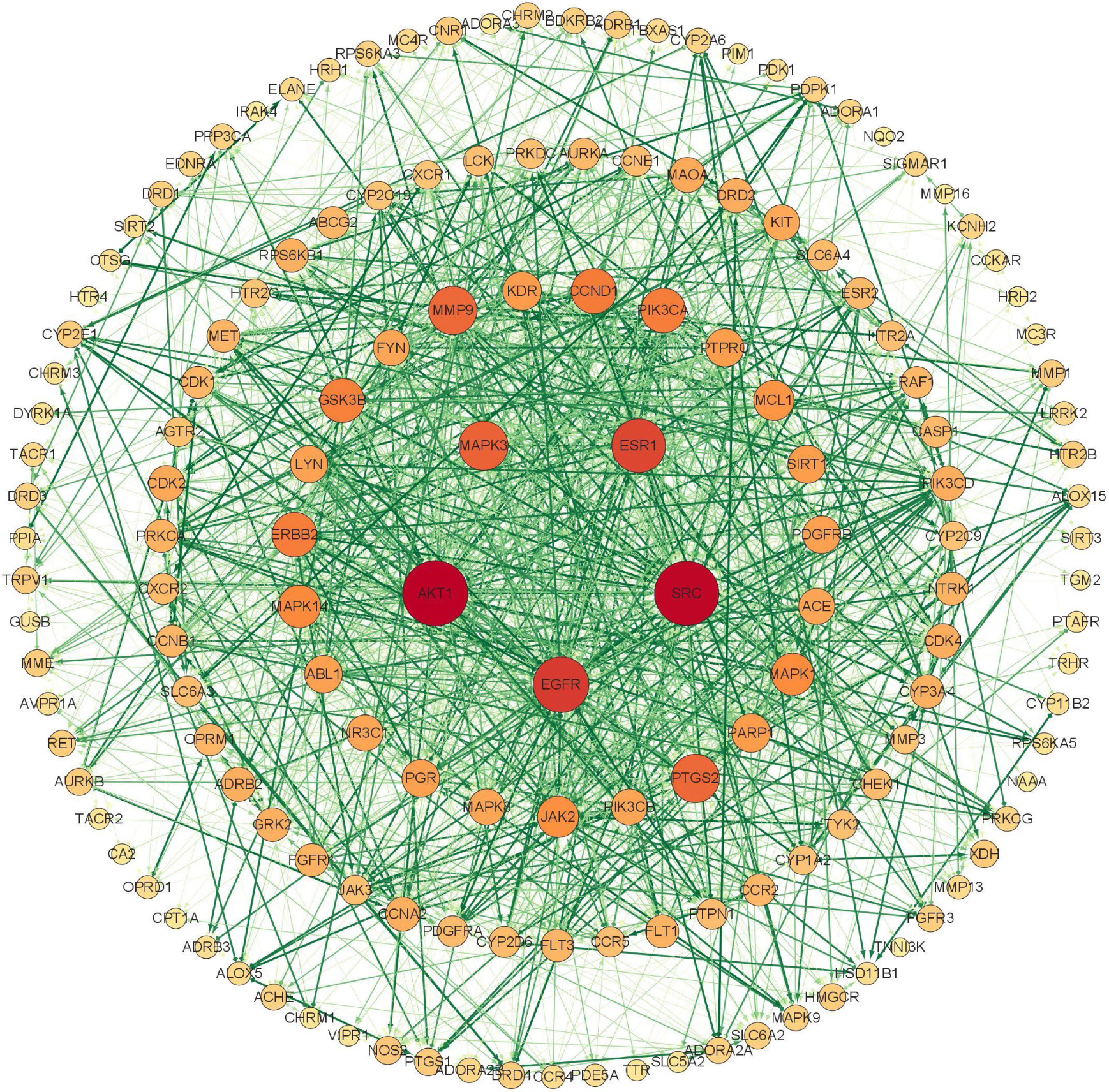
The protein-protein network interactions of potential targets.

Next, from the network analysis, we identified 23 core targets of Aflatoxin B1-induced liver toxicity (**Table 1**). Using Cytoscape 3.9.0 software, we constructed a protein-protein interaction network of these core targets (**Fig. 4**), specifically showing their interactions. It is worth noting that the top three targets, based on degree centrality, were Serine/Threonine Kinase 1 (AKT1), Proto-oncogene tyrosine-protein kinase (SRC), and Epidermal growth factor receptor (EGFR). These targets had the highest degree centrality, resulting in larger node size and a redder color in the graph. It is widely recognised in current studies that the proteins encoded by these genes play important roles in a variety of cellular functions, including regulation of cell proliferation, cell survival and cell growth, mediation of cellular immune responses, and cell signalling cascades. Through analysing the interactions in this network, it lays the foundation for potential molecular interactions in Aflatoxin B1-induced hepatotoxicity.

**Fig. 4.**
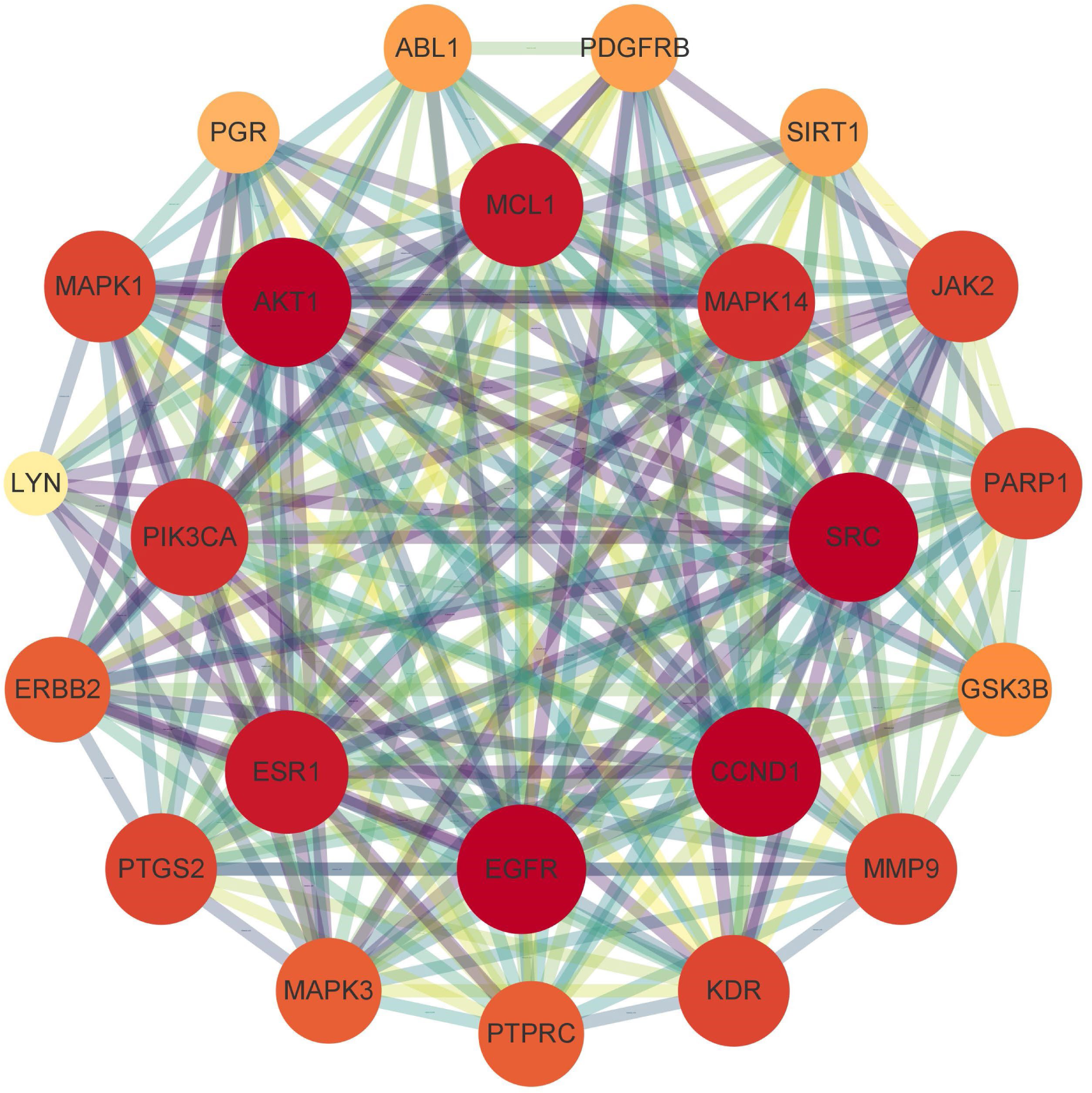
The protein-protein network interactions in core targets.

**Table 1.** Core targets screened from PPI network.

| Gene | Closeness Centrality | Degree | Radiality |
| --- | --- | --- | --- |
| AKT1 | 0.714953 | 93 | 0.995713 |
| SRC | 0.711628 | 91 | 0.995643 |
| EGFR | 0.656652 | 74 | 0.994378 |
| ESR1 | 0.634855 | 70 | 0.993815 |
| MAPK3 | 0.619433 | 61 | 0.993394 |
| PTGS2 | 0.607143 | 59 | 0.993042 |
| MMP9 | 0.602362 | 59 | 0.992902 |
| ERBB2 | 0.590734 | 52 | 0.99255 |
| GSK3B | 0.588462 | 51 | 0.99248 |
| CCND1 | 0.588462 | 53 | 0.99248 |
| MAPK14 | 0.581749 | 48 | 0.992269 |
| MAPK1 | 0.573034 | 47 | 0.991988 |
| PIK3CA | 0.570896 | 52 | 0.991918 |
| JAK2 | 0.564576 | 45 | 0.991707 |
| MCL1 | 0.564576 | 43 | 0.991707 |
| PDGFRB | 0.558394 | 38 | 0.991496 |
| SIRT1 | 0.556364 | 39 | 0.991426 |
| PGR | 0.556364 | 38 | 0.991426 |
| KDR | 0.554348 | 40 | 0.991356 |
| PARP1 | 0.554348 | 40 | 0.991356 |
| ABL1 | 0.552347 | 38 | 0.991285 |
| PTPRC | 0.552347 | 39 | 0.991285 |
| LYN | 0.534965 | 37 | 0.990653 |

### 3.4. Functional and pathway enrichment analysis of potential targets

We conducted a GO analysis of 156 potential targets using the Metascape database, restricting the species to Homo sapiens. The analysis yielded a total of 1891 statistically significant GO entries for 1591 biological processes (BP), 104 cellular components (CC) and 196 molecular functions (MF). The GO entries were sorted according to the false discovery rate (FDR) values, and we show the top 10 BP, CC, and MF with the lowest FDR values (**Fig. 5**). The Metascape database was also used to perform KEGG analysis on these 156 potential targets to determine their roles in specific signalling pathways. The top 20 KEGG signalling pathways with the lowest FDR values were visually represented in the enriched 184 signalling pathways by producing significance statistical bubble plots and category histograms (**Fig. 6**).

**Fig. 5.**
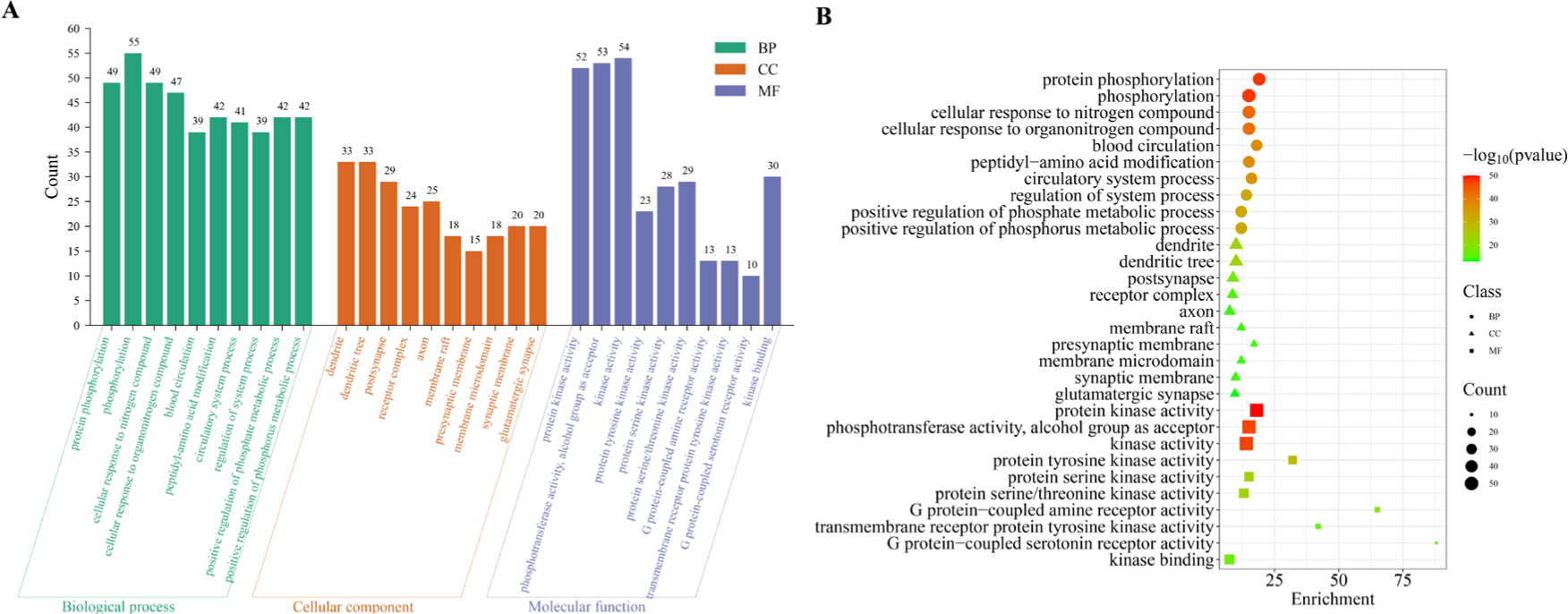
Enrichment analysis of GO for potential targets (top 10). (A)This histogram illustrated the top 10 enriched entries for each GO category (BP, CC, and MF) with smaller FDR values on the 156 potential targets. (B)The size of each bubble corresponded gene expressions in a particular pathway.

**Fig. 6.**
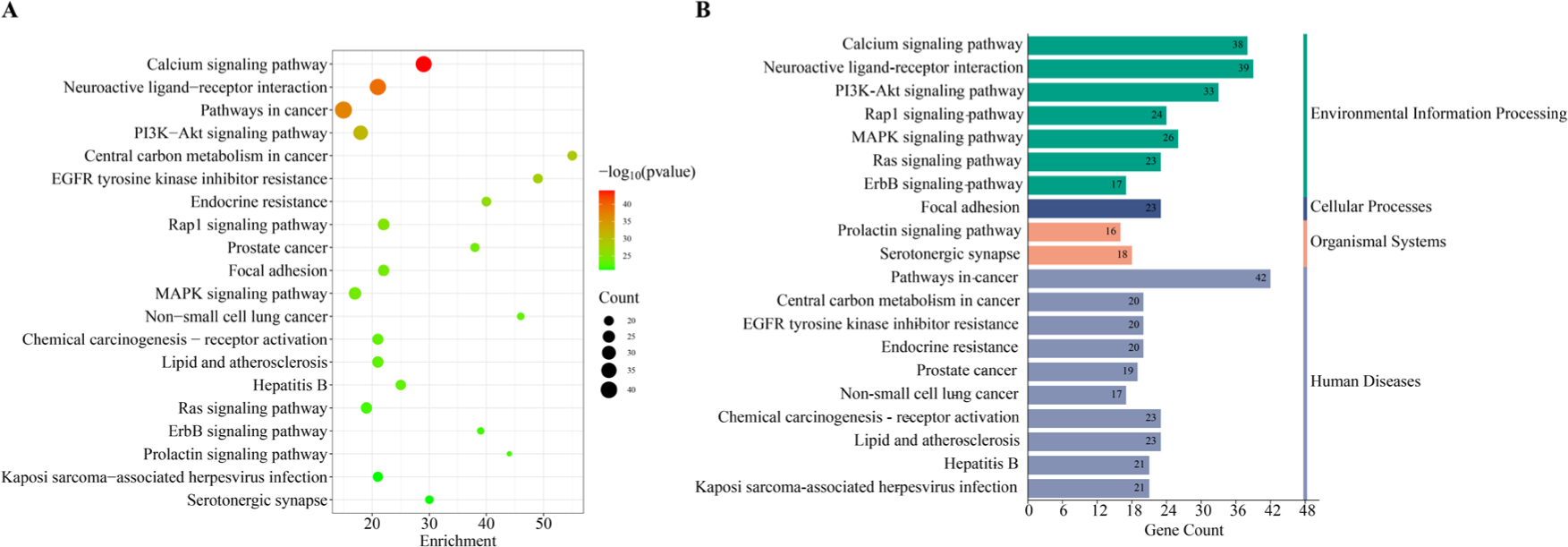
Enrichment analysis of KEGG for potential targets (top 20). (A)The bubble diagram visualized the top 20 enriched KEGG signal pathways in reverse order of FDR values. (B)The histogram illustrated the frequency and significance of enrichment for each pathway.

Notably, based on the results of GO and KEGG analyses of potential targets, these genes showed broad distribution and expression in different subcellular localisations, and many of them were involved in key regulatory processes such as protein phosphorylation, blood circulation, and circulatory system process, etc. It also functions in protein kinase activity, phosphotransferase activity, alcohol group as acceptor and kinase activity. Among the KEGG signalling pathways enriched, those associated with various signalling pathways appeared prominently, including those associated with the Calcium signaling pathway, Neuroactive ligand-receptor interaction, Pathways in cancer, and PI3K-Akt signaling pathway, and other related signalling pathways. All these findings are closely related to the type of liver injury caused by Aflatoxin B1 mentioned earlier.

### 3.5. Analysis of pathway enrichment for core targets

We performed KEGG enrichment analyses of 23 obtained core targets of Aflatoxin B1-induced hepatotoxicity and injury using the Metascape database. There were 135 signalling pathways in total that we enriched, and the top 20 pathways with smaller FDR values were selected for visualisation and analysis, depicting the core target enrichment Sankey diagram and KEGG pathway enrichment bubble diagram for each pathway (**Fig. 7**).

**Fig. 7.**
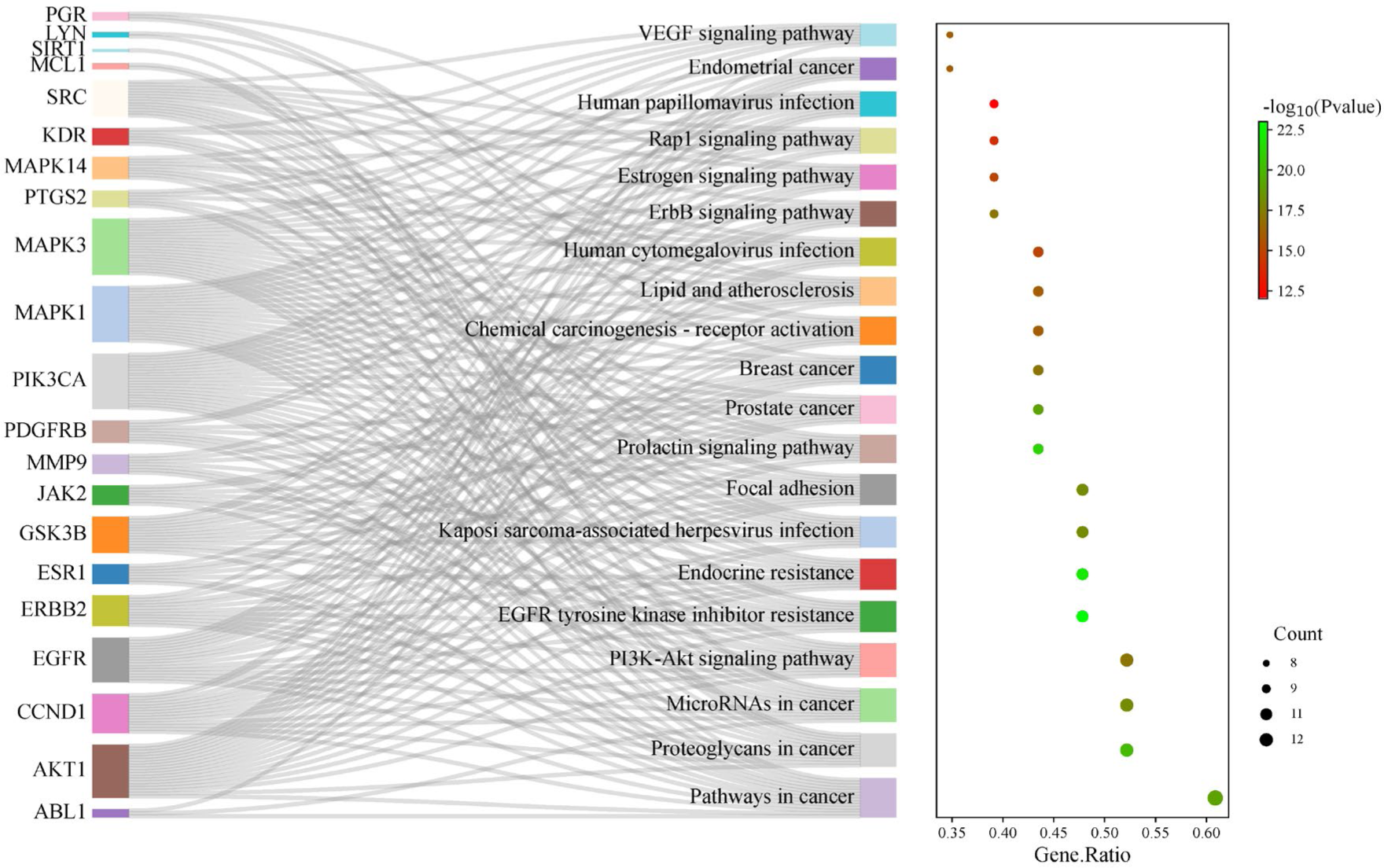
Enrichment of KEGG pathway for core targets Sankey and bubble diagrams.

The major pathways of the core targets associated with Aflatoxin B1-induced liver injury are also closely related to pathways in a variety of cancers, including prostate, breast, and endometrial cancers, as demonstrated in our study. It also involves different molecularly mediated signalling transduction, such as PI3K-Akt signaling pathway and Prolactin signaling pathway. It was interesting to note that there was a correlation between liver injury and prostate cancer, and this correlation was demonstrated for both potential and core targets (**Fig. 6**, **Fig. 7**). While further studies are needed, this finding seems to indicate potential damage to prostate tissue by Aflatoxin B1.

### 3.6. Molecular docking with core target proteins of Aflatoxin B1 to liver injury

It was performed a molecular docking analysis in order to investigate the interaction between Aflatoxin B1 and six core target proteins (AKT1, EGFR, ESR1, MAPK3, PTGS2 and SRC). Quick Vina software was used to generate six docking results quickly and showed low binding energies. These results demonstrated a strong affinity between Aflatoxin B1 and the core target proteins. Notably, the binding energies of each of the six core target proteins and Aflatoxin B1 were less than -7 kcal/mol, which indicated that Aflatoxin B1 spontaneously binds to each of the core target proteins, implying that they play an important role in the molecular mechanism of Aflatoxin B1-induced hepatotoxicity and liver injury. Furthermore, we created visualisations of the lowest binding energies between each target and Aflatoxin B1 and plotted binding energy heat maps using Discovery studio visualizer and PyMOL (**Fig. 8**, **Fig. 9**).

**Fig. 8.**
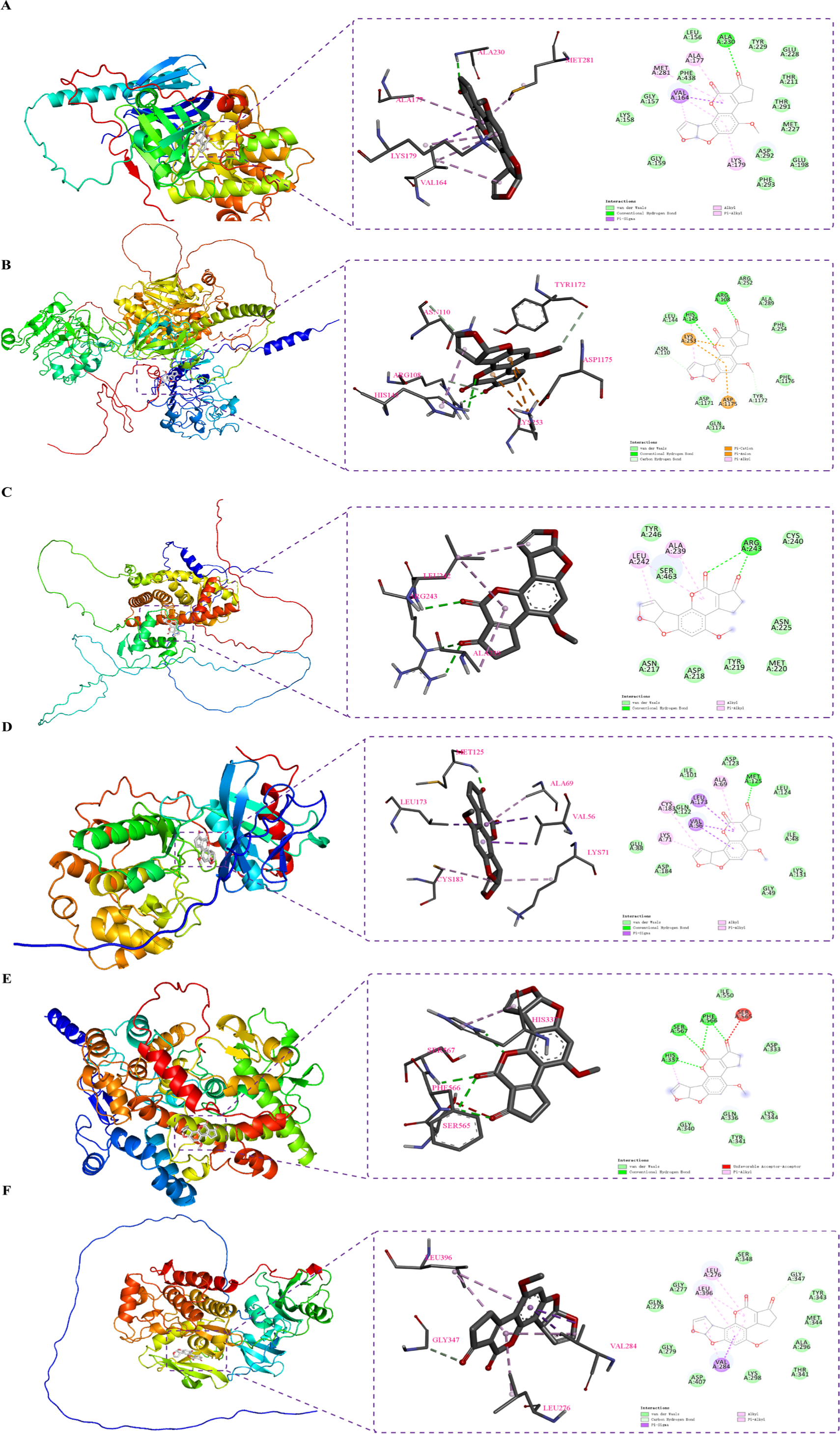
Visualisation of molecular docking results for core targets. (A) Aflatoxin B1 and AKT1, (B) Aflatoxin B1 and EGFR, (C) Aflatoxin B1 and ESR1, (D) Aflatoxin B1 and MAPK3, (E) Aflatoxin B1 and PTGS2, (F) Aflatoxin B1 and SRC.

**Fig. 9.**
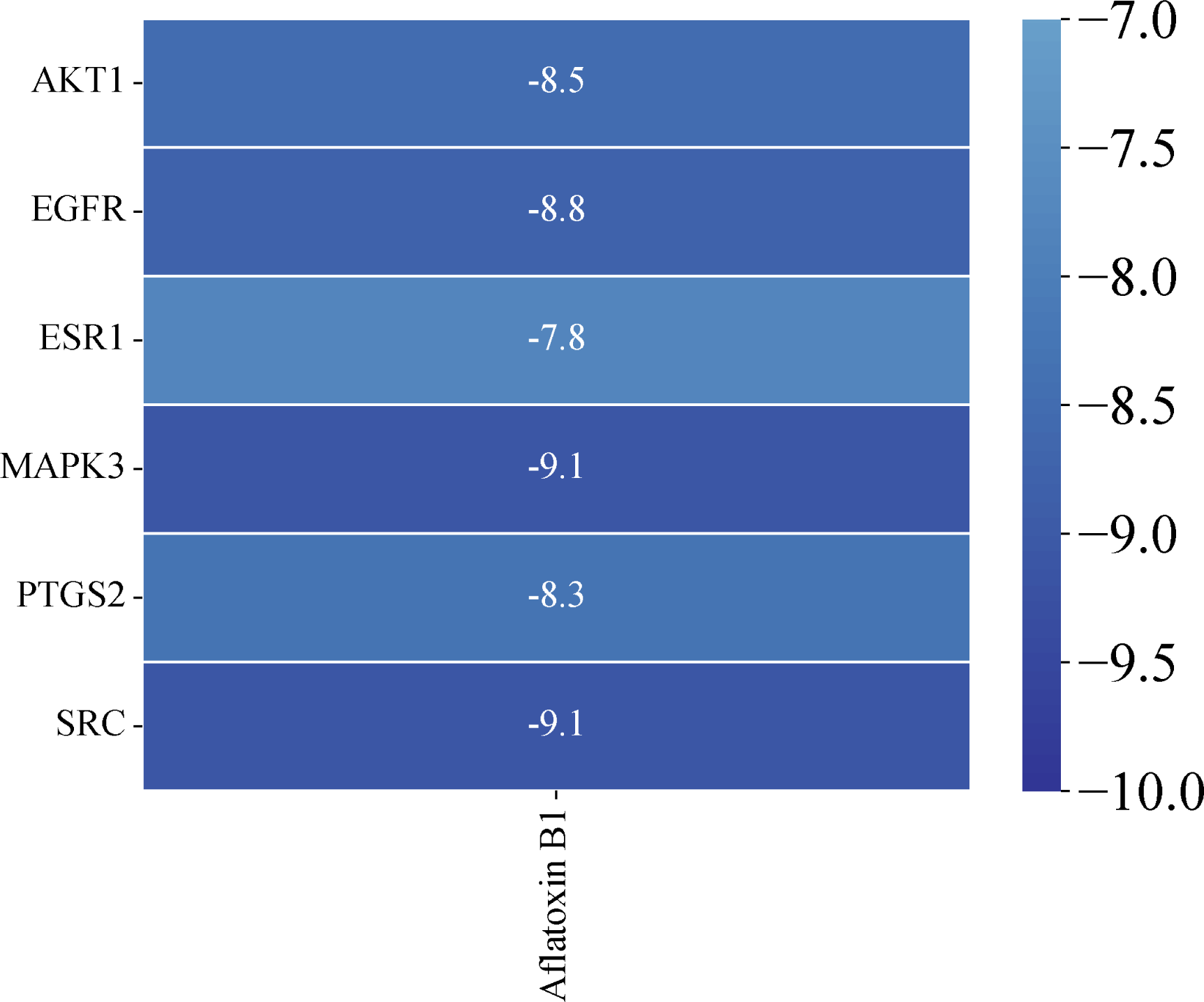
Heatmap of molecular docking binding energy of Aflatoxin B1 to core targets.

## 4. Discussion

After the initial assessment of Aflatoxin B1 toxicity in our study by first applying ProTox-II and ADMETlab 2.0 databases, we found that Aflatoxin B1 may induce hepatotoxicity. There were also 156 potential targets associated with liver injury that we screened using the ChEMBL, SwissTargetPrediction, GeneCards and DisGeNET database systems. We have established an interaction network of potential targets based on the STRING 5.0 platform and Cytoscape 3.9.0 software, and we have screened 23 core targets again, including AKT1, SRC, EGFR, ESR1, MAPK3, and PTGS2, which are the core targets of Aflatoxin B1-induced liver injury.

Previous studies have demonstrated that AKT1 plays an important role in various liver injuries, such as chronic fluorosis-induced liver injury, liver inflammation, and liver fibrosis[30, 31]. Furthermore, the analysis of our KEGG-enriched pathway also indicated that the PI3K-Akt signaling pathway involved in AKT1 also plays an important role in liver injury, such as the significantly elevated expression of PI3K and AKT1 proteins, increased apoptosis, and increased intracellular calcium concentration in hepatocytes with fluorosis[30]. Hence, we suggested that AKT1 and its PI3K-Akt signaling pathway also play an important role in liver injury.

The SRC, which is encodes a membrane-associated non-receptor tyrosine kinase, has an essential role in cell growth, division, migration and survival signalling pathways[32]. Firstly, Src kinase can also regulate glucose metabolism in cancer cells through different mechanisms[33]. Second, it has been shown that SRC plays an important role in endotoxin-mediated liver injury[34]. The above studies are consistent with our results, which suggested that SRC plays an important role in liver injury and cancer development.

Hepatocyte-specific EGFR activity promotes cholestatic injury and activates pro-inflammatory responses, which promote cholestatic liver disease[35]. Alcoholic hepatitis is a serious liver injury phenomenon, and it has been demonstrated that the E3 ubiquitin ligase NEDD4 promotes EGFR internalisation in alcoholic hepatitis, thereby affecting related signalling pathways to promote liver regeneration[36]. Moreover, activation of EGFR triggers cellular autophagy, and Schisandrin B reduces the expression level of EGFR to mitigate the harmful effects of liver injury[37]. Hence, EGFR plays an essential role in liver injury.

In vivo experiments demonstrated that high expression of ESR1 leads to apoptosis and inflammation in hepatocytes, and that ERBB2 expression can be enhanced by suppressing ESR1 expression to attenuate hepatocyte apoptosis and inflammatory responses[38]. Besides, ESR1-mediated signalling inhibited liver regeneration by down-regulating Wnt signalling, which led to reduced activation of cell cycle protein D1 after chemical-induced liver injury[39]. Consequently, ESR1 has an important role in Aflatoxin B1-induced liver injury.

MAPK3 plays a mediating role in the pathogenesis, progression, metastasis, drug resistance and poor prognosis of a variety of malignant tumours, such as gliomas, hepatocellular carcinomas and ovarian cancers[40]. We discovered that MAPK3 was involved in multiple signalling pathways, including Pathways in cancer, Proteoglycans in cancer, MicroRNAs in cancer, and other multiple cancer signalling pathways by KEGG enrichment analysis. Therefore, it was hypothesised that MAPK3 plays an important role in Aflatoxin B1-induced liver injury.

There are studies that have discovered that in colorectal cancer patients, PTGS2 expression is associated with an enhanced risk of tumour recurrence and a decreased colorectal cancer-specific survival rate[41]. The participation of PTGS2 in multiple signalling pathways such as Pathways in cancer, VEGF signaling pathway, and Human papillomavirus infection was discovered in our research. It is hypothesised that Aflatoxin B1 can induce liver injury or hepatocellular carcinoma by affecting the expression of PTGS2, which regulates multiple signalling pathways such as cancer.

The functional analysis of potential and core targets and KEGG enrichment analysis of Aflatoxin B1-induced liver injury demonstrated that Aflatoxin B1 affects multiple targets and pathways to induce liver injury. This result is consistent with the current study. Furthermore, it is illustrated in published studies that exposure to Aflatoxin B1 induces liver injury, hepatocellular carcinoma, and many other cancers[11, 13]. This result is enough to prove that our research method is accurate and reliable.

The toxicity of Aflatoxin B1 was initially analysed by us through literature research and ProTox-II and ADMETlab 2.0. Then, we have collected 156 potential targets of Aflatoxin B1 induced liver injury by querying using ChEMBL, SwissTargetPrediction, GeneCards and DisGeNET databases. Subsequently, we have constructed a protein-protein interaction network by screening 23 core targets including AKT1, SRC, EGFR, ESR1, MAPK3 and PTGS2 using STRING5.0 database and Cytoscape 3.9.0. Notably, these core targets play an essential role in tumourigenesis. Lastly, based on the Metascape database, we performed GO enrichment analysis and KEGG pathway enrichment analysis for potential and core targets. The results of our study has revealed the significance of signalling pathways associated with liver injury and hepatocellular carcinoma, such as Pathways in cancer, PI3K-Akt signaling pathway, Rap1 signaling pathway, and other pathways, and the network regulation between these signaling pathways is closely related to liver injury. It indicated that Aflatoxin B1 could induce the occurrence of liver injury, and at the same time promote the apoptosis of hepatocytes, which eventually led to liver dysfunction. The molecular docking results demonstrated that all the six core targets were stably bound to Aflatoxin B1, and the binding energies were lower than -7 kcal/mol, suggested that the six core targets play an important role in Aflatoxin B1-induced liver injury.

The current study, apart from providing a molecular mechanism for the hepatotoxicity caused by Aflatoxin B1, proposes the use of network toxicology and molecular docking strategies for the rapid study of the toxicity of chemical contaminants during food production, processing and preservation. Compared with traditional food toxicological evaluations, we evaluated the toxicity of chemical substances in food contaminants through network toxicology and molecular docking, which effectively saves time and resources, enables large-scale data processing, integrates multiple influencing factors, enables the exploration of mechanisms and the prediction of new toxicities, as well as reduces the need for and avoids ethical issues in animal experiments.

In addition, network toxicology can integrate multiple influencing factors, such as chemical structure, biological activity and metabolic pathways, as well as epidemiological information based on big data[3, 4]. Meanwhile, network toxicology explores the interactions between chemical substances and molecular and cellular mechanisms in living organisms through methods such as systems biology and network analyses, thereby predicting new toxic effects[3, 4]. It helps to identify potential toxicity risks in advance. Molecular docking technique that is used to predict the binding mode and affinity between small molecules and proteins or other biological macromolecules[6, 27]. The use of network toxicology and molecular docking techniques can efficiently and accurately assess the toxicity of chemicals in food contaminants, providing effective protection for food safety and people’s health. Network toxicology and molecular docking techniques should be used in the future assessment of food contaminant toxicity for large-scale data analysis, which can be combined with animal models if necessary to confirm the core targets and core signalling pathways of Aflatoxin B1-induced liver injury, and use this as a breakthrough point to search for remedies for the prevention and treatment of Aflatoxin B1-induced liver injury.

## 5. Conclusions

In conclusion, we have provided a comprehensive analysis of the potential hepatotoxicity of Aflatoxin B1 in our study through the strategies of network toxicology and rapid molecular docking analysis. There were 156 potential targets of Aflatoxin B1-induced liver injury, and we further screened 23 core targets, such as AKT1, SRC and EGFR, which may play important roles in mediating the toxic effects of Aflatoxin B1 on the liver. As a result of KEGG enrichment analysis we enriched several important signalling pathways, such as Cancer-related signalling pathway, PI3K-Akt signaling pathway, Chemical carcinogenesis - receptor activation and so on. The results of our study suggested that exposure to Aflatoxin B1 may lead to liver injury and increase the risk of carcinogenesis through modulation of cell proliferation, cell survival, cell growth, cellular immune response, and cellular signalling cascade responses in hepatocellular carcinoma cells.

The present study not only elucidates the potential molecular mechanisms of hepatotoxicity of Aflatoxin B1, but also promotes network toxicology and molecular docking strategies to effectively analyse the toxicity and molecular biological mechanisms of potential food contaminants, as well as effectively addressing the limitations of time, cost, ethical issues of animal use, and predictive power of traditional toxicological assessments, and lays the foundation for research on diagnosis of diseases associated with exposure to these toxic substances.

## CRediT authorship contribution statement

Zi-Yong Chu designed the experiment and performed the statistical analysis. Zi-Yong Chu and Xue-Jiao Zi carried out the study. Zi-Yong Chu and Xue-Jiao Zi drafted the manuscript. All authors read and approved the final manuscript.

## Acknowledgement

The authors declare that they have no known competing financial interests or personal relationships that could have appeared to influence the work reported in this paper. This research did not receive any specific grant from funding agencies in the public, commercial, or not-for-profit sectors.

## Data availability

The data that has been used is confidential.

